# Structural diversity and distribution of NMCP-class nuclear lamina proteins in streptophytic algae

**DOI:** 10.1101/2024.07.24.604981

**Authors:** Brendan S. Kosztyo, Eric J. Richards

## Abstract

Nuclear Matrix Constituent Proteins (NMCPs) in plants function like animal lamins, providing the structural foundation of the nuclear lamina and regulating nuclear organization and morphology. Although they are well-characterized in angiosperms, the presence and structure of NMCPs in more distantly related species, such as streptophytic algae, are relatively unknown. The rapid evolution of NMCPs throughout the plant lineage has caused a divergence in protein sequence that makes similarity-based searches less effective. Structural features are more likely to be conserved compared to primary amino acid sequence; therefore, we developed a filtration protocol to search for diverged NMCPs based on four physical characteristics: intrinsically disordered content, isoelectric point, number of amino acids, and the presence of a central coiled-coil domain. By setting parameters to recognize the properties of bona fide NMCP proteins in angiosperms, we filtered eight complete proteomes from streptophytic algae species and identified strong NMCP candidates in six taxa in the Classes Zygnematophyceae, Charophyceae, and Klebsormidophyceae. Through analysis of these proteins, we observed structural variance in domain size between NMCPs in algae and land plants, as well as a single block of amino acid conservation. Our analysis indicates that NMCPs are absent in the Mesostigmatophyceae. The presence versus absence of NMCP proteins does not correlate with the distribution of different forms of mitosis (e.g., closed/semi-closed/open) but does correspond to the transition from unicellularity to multicellularity in the streptophytic algae, suggesting that an NMCP-based nucleoskeleton plays important roles in supporting cell-to-cell interactions.

**SIGNIFICANCE STATEMENT:** All eukaryotic organisms contain a membrane-bound nucleus, which holds a cell’s DNA, but plants and animals have distinct sets of proteins that make up the mesh-like scaffold inside the nucleus (i.e., nuclear lamina) that is essential for structural integrity and organization. Nonetheless, the major nuclear lamina proteins in plants and animals share structural features, and we exploited these characteristics to identify a key class of nuclear lamina proteins in streptophytic algae, allowing us to chart the distribution of these proteins across photosynthetic organisms and to gain insight into the evolution of nuclear organization. Our results indicate that the major class of nuclear lamina protein in plants evolved independently from structurally similar nuclear lamina proteins in animals.

## INTRODUCTION

In eukaryotes, the nucleus is central to the integration and execution of cell functions, including storing the genetic material and regulating its expression. A complex of proteins called the nuclear lamina (NL), which resides directly below the inner nuclear membrane, is responsible for the organization of the nucleus and its indirect attachment to the cytoskeleton, as well as for genome and transcription factor regulation (Gruenbaum and Foisner, 2015). In metazoans, lamins are the major structural protein of the NL and the founding member of the intermediate filament superfamily (Dechat *et al*., 2010).

Despite being ancestral and well conserved, lamins are not found constitutively throughout the eukaryotic lineage. The absence of lamin homologs could be explained by an extreme divergence in certain eukaryotic lineages (Dittmer and Misteli, 2011). Alternatively, the NL could have independent origins, being composed of distinct proteins recruited to form the NL through convergent evolution (Koreny and Field, 2016). Distinguishing between these two hypotheses requires assembly of a comprehensive catalog of lamin homologs and potential functional analogs across divergent eukaryotic lineages.

In the non-metazoan *Dictyostelium discoideum*, NE-81 represents an evolutionary precursor to the A- and B-type lamins found in vertebrates (Batsios *et al*., 2012; Odell and Lammerding, 2023). This *Dicytostelium* protein has the structural features of metazoan lamins, with which it shares primary amino acid sequence similarity (Krüger *et al*., 2012). Further, transgenic expression of NE-81 in a lamin A/C deficient cell line partially complements the mechanical defects exhibited by nuclei in those cells (Odell *et al*., 2023). In other non-metazoan groups, distinct NL proteins exist that are difficult to reconcile as lamin homologs. For example, in Trypanosomes, NUP-1 and NUP-2 are NL proteins that appear to function analogously to lamins, but these proteins lack sequence similarity with lamins and share only some structural features (e.g., long coiled-coil domains in NUP-1; see Figure 1) (DuBois *et al*., 2012; Maishman *et al*., 2016; Padilla-Mejia *et al*., 2021).

**Figure 1:**
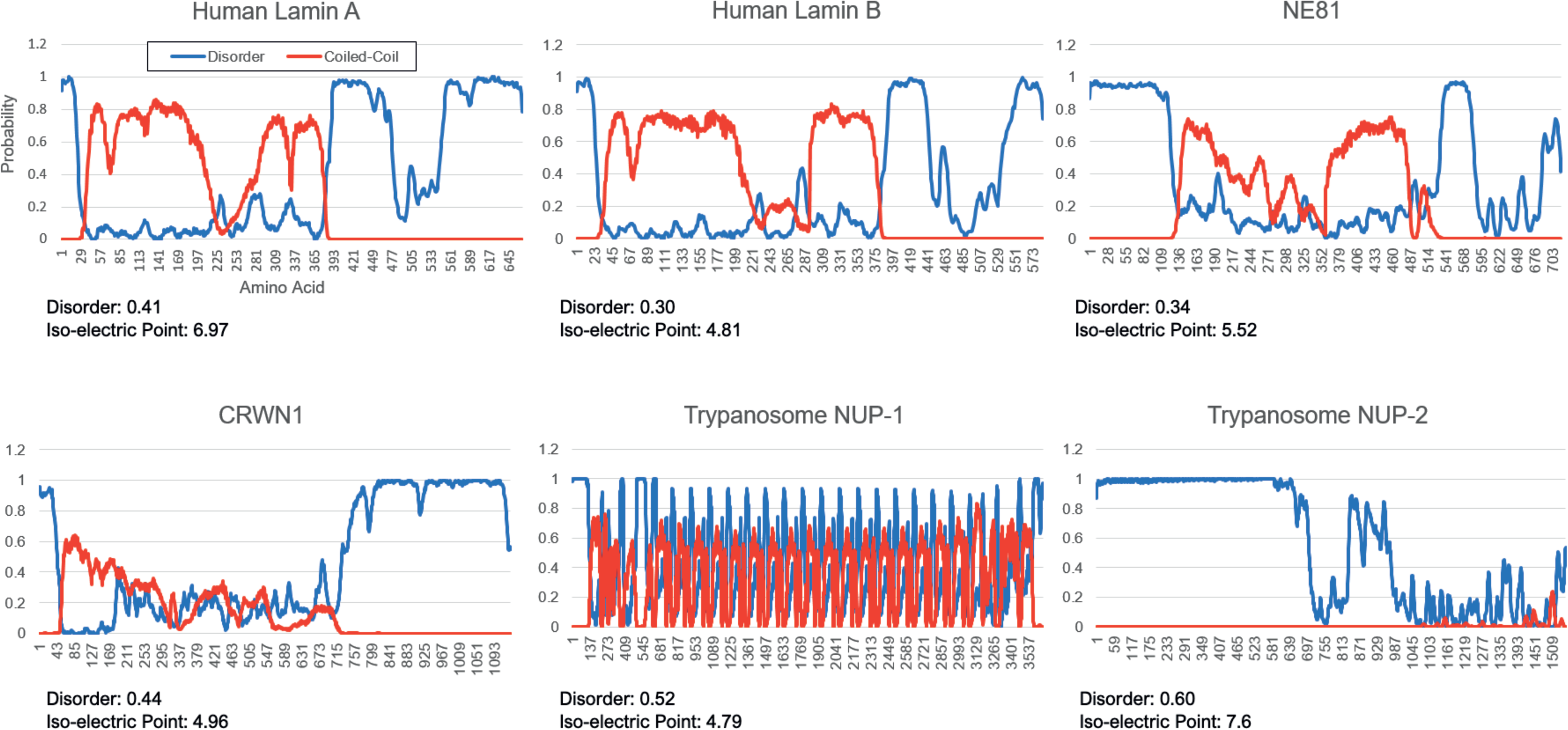
Structural comparison of NL proteins in different eukaryotes. Scores for intrinsically disordered (blue) and coiled-coil (red) regions were mapped per residue for human lamin A, human lamin B1, Trypanosome NUP-1 and -2, NE81 in *Dictyostelium*, and CRWN1 in *Arabidopsis*. Human lamins, CRWN1, and NE81 share an overall tripartite organization, with disordered regions flanking a long, central coiled-coil region. NE81 differs from human lamins and CRWN proteins in having an elongated N-terminus. The Trypanosome proteins lack this tripartite organization. Note that the proteins lengths are normalized and not drawn to scale.

The situation in plants is more complex. Plants lack recognizable lamins yet nuclei in plant cells have a NL structure (Masuda *et al*., 1993; Fiserova *et al*., 2009). The major component of this structure in plant cells are Nuclear Matrix Constituent Proteins (NMCPs), first discovered in the angiosperm *Daucus carota* (carrots) (Masuda *et al*., 1997; Kimura *et al*., 2010; Ciska and Moreno Díaz de la Espina, 2013). Functional studies in *Arabidopsis thaliana*, demonstrate that NMCPs (also called CRWNs, for CROWED NUCLEI, in this species) are involved in the regulation of nuclear size and shape as well as heterochromatin organization (Dittmer *et al*., 2007; Sakamoto and Takagi, 2013; Wang *et al*., 2013; Sakamoto *et al*., 2022). NMCPs in angiosperms are essential for survival (Wang2013; Blunt *et al*., 2023), but the sole *NMCP* gene can be deleted without affecting viability in the liverwort *Marchantia polymorpha*, despite the fact that disruption of three-dimensional chromatin organization results (Wang *et al*., 2021). These functional studies indicate that NMCPs play similar roles to lamins, but these major NL proteins in plants do not share significant amino acid sequence similarity with lamins. Nor do they contain diagnostic features of lamins, such as a C-terminal CAAX box or an immunoglobulin-fold domain (Ciska *et al*., 2019). Nonetheless, NMCPs and lamins share structural characteristics, as illustrated in Figure 1. The most obvious similarity is that NMCPs and lamins have a tripartite structure defined by a long central coiled-coil domain. Another feature shared by NMCPs and lamins is the high percentage of intrinsically disordered regions (IDRs) in the N- and C-terminal domains (Huang *et al*., 2021; Reimann *et al*., 2023; Qin *et al*., 2011). These structural similarities suggest that NMCPs could be highly diverged lamin homologs, but these similarities are also consistent with the convergent evolution hypothesis.

To determine if NMCPs are distant lamin homologs or evolutionarily distinct, we broadened our search for NMCPs in the ‘green lineage,’ encompassing land plants and photosynthetic algae (Merchant *et al*., 2007). All land plants examined to date contain NMCPs, including angiosperms, gymnosperms, ferns, liverworts and mosses (Poulet *et al*., 2016; Ciska *et al*., 2019). Recent similarity-based studies have shown that the presence of NMCPs in the green lineage extends as far as streptophytic algae, a sister group to land plants, but obvious NMCPs are absent in more distantly related green algae (Chlorophyta) (Koreny and Field, 2016; Ciska *et al*., 2019).

Recognizable NMCPs are extremely divergent, suggesting that the primary amino acid sequence similarity might not be an effective approach to find ancestral orthologs (Ciska *et al*., 2019). Moreover, unlike mammalian lamins, NMCPs lack conserved domains, such as diagnostic post-translational modification sites, a C-terminal CaaX prenylation motif, and an immunoglobulin domain, making sequenced-based identification even more challenging (Ciska and Moreno Díaz de la Espina, 2013). We reasoned that overall structural organization would more likely be conserved and therefore more useful in searching for distantly related proteins. Accordingly, we designed a method of proteomic filtration using protein properties rather than amino acid sequence similarity to search for highly diverged NMCP orthologs in the green lineage. One criterion involved filtering proteins based on polypeptide size (> 800 amino acids) and acidic isoelectric point (IEP, pH 4.0 to 6.0). A second criterion was the presence of an extended coiled-coil domain. A third critical criterion of our filtration protocol was a high incidence of intrinsically disordered regions (IDRs), diagnostic of all characterized NMCPs and also found in animal lamins (see Materials and Methods). Intrinsically disordered regions (IDRs) are domains within proteins that lack a defined three-dimensional shape (Piovesan *et al*., 2022). One possibility is that having a malleable shape allows for plasticity in binding, not attainable by ordered proteins, and dynamic structural ordering (Nishiyama *et al*., 2018). Phase separation is another emerging theory for IDR function, in which cellular components transition between fluid and more rigid states of matter, often occupying a medial state (Molliex *et al*., 2015). IDRs promote these phase transitions by acting as linkers and stimulating sticky interactions with other proteins, mediated by weak hydrophobic interactions (Banjade *et al*., 2015; Boeynaems *et al*., 2018). Phase separation and condensate formation from such interactions among proteins could contribute to these proteins’ organization of the nuclear lamina.

We focused our search on well-characterized genomes and deduced proteomes of streptophytic algae, which comprise diverse photosynthetic organisms in sister lineages to land plants, with the aim of understanding the evolutionary history of the main component of the plant NL. Our results indicate that NMCP-class NL proteins are restricted to a subset of streptophytic algae, and we found no evidence for transitional states with features more closely related to animal lamins. These findings support the view that organisms in the plant kingdom contain a convergently evolved NL composed of diverse NMCP proteins, rather than specialized plant-specific lamin orthologs.

## RESULTS

We began our study by selecting sequenced genomes from eight different taxa in four different Classes of streptophytic algae (see Figure 2). In addition, we examined one green algal (chlorophytic) species (*Chlamydomonas reinhardtii*), one moss species (*Physcomitrium patens*), and one angiosperm species (*Arabidopsis thaliana*) as reference taxa bracketing the streptophytic algae.

**Figure 2:**
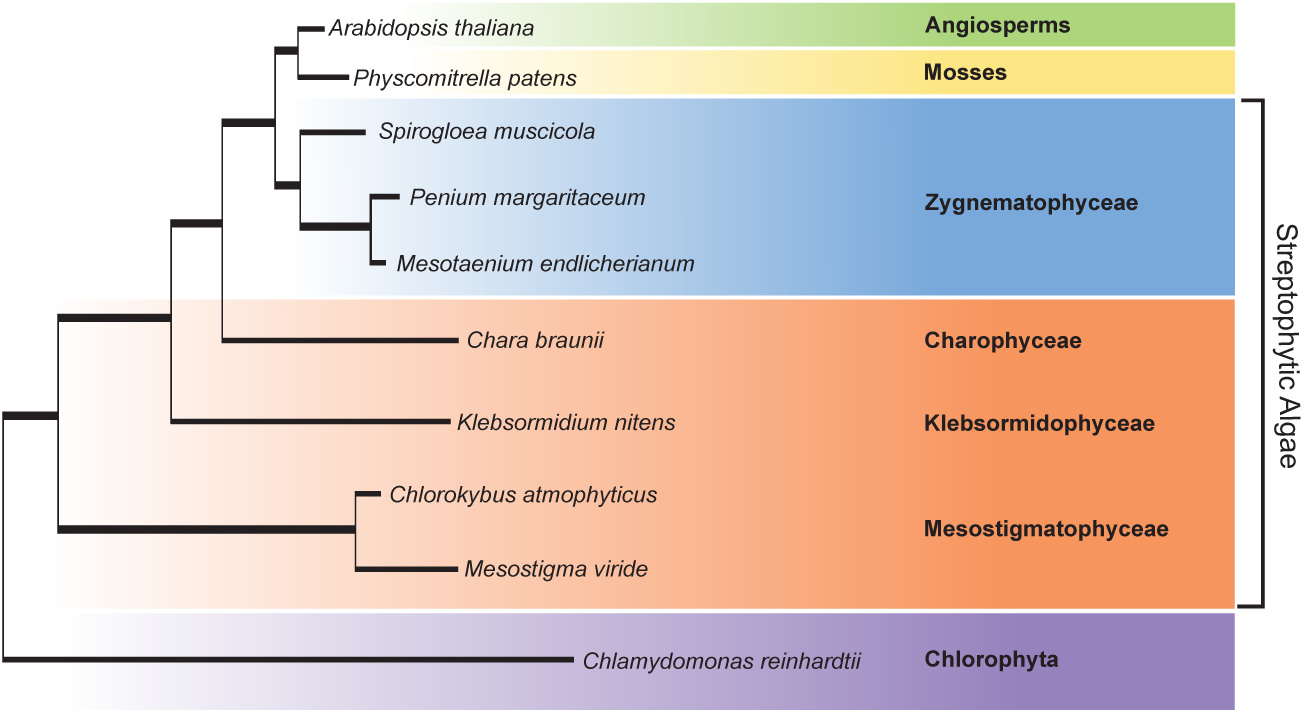
Phylogenetic relationships among species that were used in this study. Branch lengths are proportional to evolutionary distance, based on Figure 1 from Cheng *et al*. (2019). The modified position of Spirogloea between land plants and Zygnematophyceae is supported by the recent work of Hess *et al*. (2022). Four Classes of streptophytic algae were examined: Zygnematophyceae, Charophyceae, Klebsormidophyceae, and Mesostigmatophyceae. Charophyceae, Klebsormidophyceae, and Mesostigmatophyceae are closely related and grouped together (in orange) because of similarity in their morphological characteristics. Green algae (Chlorophyta) are more diverged from land plants (represented here by Angiosperms and Mosses) than streptophytic algae.

Deduced proteomes from the sequenced genomes were processed following the filtration steps diagrammed in Figure 3. The first characteristic used to filter each proteome was the presence of intrinsically disordered regions (IDRs), which are diagnostic of NMCP and lamin proteins (see Materials and Methods). We use the program Metapredict to locate regions predicted to have high disordered content and to filter for protein size. NMCP proteins are typically very large; in all known land plants NMCPs are ≥1000 amino acids, therefore, we selected a cutoff of 800 amino acids for filtration. Independently, we used the program Emboss to filter for protein isoelectric point (IEP). Known NMCPs have an acidic IEP, around 5.3. We chose a threshold of IEP between 4-6 to filter the deduced proteomes. The results from these three steps were collected, and the tool Venny: Venn Diagrams (https://bioinfogp.cnb.csic.es/tools/venny/) was used to select proteins that satisfied all three filters.

**Figure 3:**
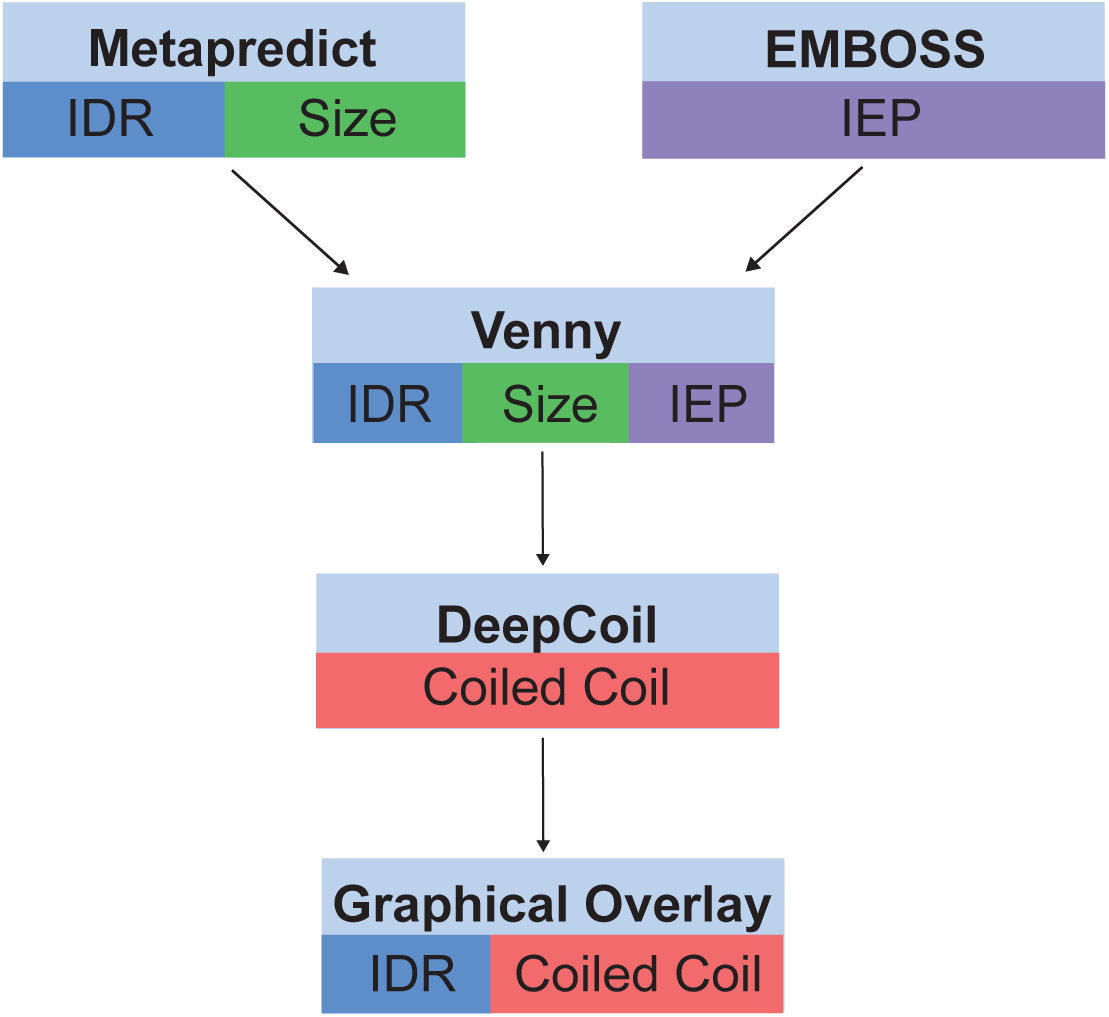
Summary of proteome filtration pathway. Each filtration step is represented by a separate box. The method used to execute each filtration step is shown in the upper half of each box, and the characteristic being evaluated is shown in the lower portion. Intrinsic disorder is represented in blue, size is represented in green, isoelectric point is shown in purple, and coiled-coil is shown in red.

With the condensed list, we used the program DeepCoil to generate the probability of finding a coiled-coil domain at each amino acid. These results were expressed graphically and analyzed visually for similarity to the NMCP coiled-coil region. The proteins selected had a central coiled-coil region that spanned at least 100 amino acids. We then overlaid IDR scores on coiled-coil maps for each selected protein, and the composite graphical representations were visually inspected to generate a list of finalists.

We applied this filtration protocol to the *A. thaliana* proteome and recovered seven protein “finalists,” four of which were bona fide NMCPs (i.e., CRWN1-4). The remaining three proteins are predicted to be filament-like proteins, including TCS1 (TRICHOME CELL SHAPE 1), which is a cortical microtubule binding protein (Chen *et al*., 2016). Two additional proteins of unknown function were found (At1g47900 and the SMAD/FHA domain protein encoded by *At2g45460*). In the moss, *Physcomitrium patens*, three finalists were identified by the filtration protocol, two of which are proteins that show a high degree of amino acid similarity with *A. thaliana* NMCPs. These two proteins were previously identified as NMCP homologs in *P. patens* (Wang *et al*., 2013; Ciska *et al*., 2019). The third protein (XP_024361755) shows a high degree of similarity with Arabidopsis TCS1 and is a likely moss homolog of this protein. These findings suggest that the filtration protocol is robust and identifies known NMCPs in land plant proteomes, but also retrieves a small number of additional proteins that are likely false positives not involved in nuclear lamina formation.

Consistent with this expectation, the filtration protocol identified six proteins from *Chlamydomonas reinhardtii,* a member of the green algae family, that lacks known NMCPs and was used as a negative control. Four of these proteins are predicted to have roles outside of the nucleus, including a kinesin motor domain-containing protein (A0A2K3CRT1), a tropomyosin (A0A2K3DS11), a centrosomal protein (A0A2K3D6E5), and a protein implicated in Golgi vesicle trafficking (A0A2K3D1Z8). The remaining two Chlamydomonas proteins identified by our filter did not have a predicted or characterized function.

After confirming that the filtration protocol can identify NMCPs from whole proteomes with a modest number of false positives, eight streptophytic algal species were analyzed using the same approach. Once all four physical characteristics (size, disorder (IDR), IEP, and coiled-coil region similarity) were filtered, finalists were analyzed to examine their structural organization. As described above, proteins with confirmed coiled-coil domains were put in a graphical overlay that combined coiled-coil and IDR prediction on a per-residue basis. Graphical analysis allowed for the protein to be visualized holistically and compared to angiosperm NMCP, to assess organizational similarity.

Each streptophytic algal species filtered returned between 1 and 13 finalist proteins after this graphical analysis. We know from the filtration of the control proteomes that a few false positives can be expected. Therefore, another step was needed to differentiate NMCP candidate proteins from false positives. To supplement our sequence-agnostic approach, we applied multiple sequence alignments to identify amino acid motifs that distinguished among the finalists generated by the filtration protocol. As shown in Figure 4, a single protein from five of the eight streptophytic algae species examined contained a 50-amino acid region of similarity in the coiled-coil region (see the rightmost column of Table 1). The region corresponds to a short, degenerate but conserved region in the N-terminus of the coiled-coil region previously identified in angiosperm NMCPs (Masuda *et al*., 1997; Dittmer *et al*., 2007).

**Figure 4:**
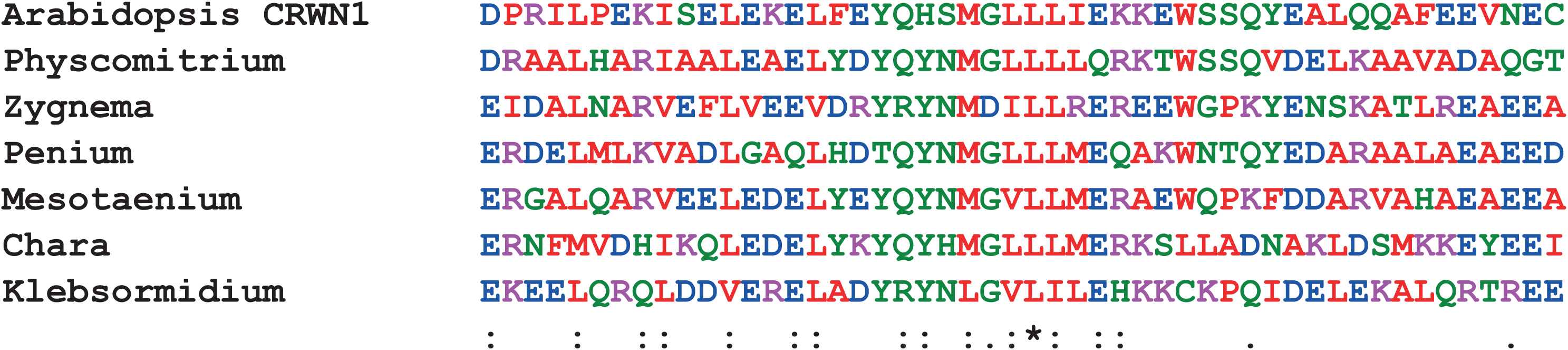
Multiple sequence alignment of a weakly conserved block within algal NMCPs. The 50 amino acids of the N-terminus of the coiled-coil regions of NMCP-like candidates and two reference NMCPs are shown. The core of the motif is a well-conserved MGLL sequence, which lies close to the N-terminal border of the extended coiled-coil domain of the proteins. Amino acids in green are polar, red are non-polar, blue are negatively charged, and pink are positively charged. The *Physcomitrium* protein, NMCP1, used in the multiple sequence alignment is A0A2K1L4A2 encoded by the *PHYPA_003649* gene.

**Table 1:** Stepwise protein count through the filtration process. This table summarizes the protein number present after each filtration step was executed. *Proteome* describes the total number of proteins before filtration. *Disorder* corresponds to the proteins satisfying the Metapredict threshold of 0.3. *Size* shows the number of proteins over 800 amino acids in length. IEP represents the proteins that had an isoelectric point in the range of 4-6 pKi. *Size, Disorder, IEP* describes the number of proteins that satisfy all three requirements. *Coiled-Coil* shows the number of proteins with a structurally similar coiled-coil region via DeepCoil (EMBL) per residue predictions, of the proteins that were included in SDI. *Filtered Finalists* describes the number of proteins, of those that had satisfied the previous requirements, with a similar graphical distribution of disorder per residue, which also have protein duplicates removed to describe the number of discrete proteins considered. *Candidates* combines the data from the coiled-coil analysis, mapping the per residue disorder scores with the per residue coiled-coil pattern to look for graphical similarities to a typical NMCP map and ensure both motifs are accounted for and is compared with a multiple sequence alignment to known NMCPs.

| Species | Proteome | Disorder | Size | IEP | Size<br>Disorder<br>IEP | Coiled<br>Coil<br>(SDI) | Filtered<br>Finalists | NMCP<br>Candidates |
| --- | --- | --- | --- | --- | --- | --- | --- | --- |
| *Arabidopsis thaliana* | 48266 | 21874 | 3539 | 12777 | 596 | 21 | 7 | 4 |
| *Physcomitrium patens* | 48022 | 28682 | 8900 | 12134 | 1985 | 19 | 3 | 2 |
| *Spirogloea muscicola* | 27137 | 15841 | 3549 | 7006 | 640 | 42 | 13 | 0 |
| *Penium margaritaceum* | 52333 | 36805 | 2094 | 13075 | 612 | 14 | 1 | 1 |
| *Mesotaenium endlicherianum* | 44092 | 28972 | 5553 | 10310 | 1128 | 29 | 8 | 1 |
| *Zygnema circumcarinatum* | 34700 | 24266 | 4299 | 7532 | 1044 | 61 | 5 | 1 |
| *Chara braunii* | 35672 | 21605 | 8157 | 11176 | 2223 | 19 | 5 | 1 |
| *Klebsormidium nitens* | 16283 | 7846 | 1305 | 4957 | 544 | 128 | 10 | 1 |
| *Chlorokybus melkonianii* | 8693 | 4339 | 851 | 2168 | 125 | 9 | 7 | 0 |
| *Mesostigma viride* | 16232 | 8231 | 3045 | 4090 | 566 | 49 | 11 | 0 |
| *Chlamydomonas reinhardtii* | 19526 | 12939 | 5051 | 4440 | 1033 | 46 | 6 | 0 |

After identifying this region of sequence similarity in the coiled-coil domain, we wanted to determine if the filtration protocol had missed any other NMCP candidates containing this sequence. The 50-amino acid region of the *P. patens* NMCP protein shown in Figure 4 was used as an independent query in a series of local BLAST searches against the proteome of each algal species. The candidate proteins observed in *M. endlicherianum, C. braunii, P. margaritaceum, Z.circumcarinatum* and *K. nitens* returned as significant matches (see Supplementary Table 1). In *P. margaritaceum,* a second protein was identified (pm015243g0010), which was annotated as a putative NMCP in the existing genome annotation (Jiao *et al*., 2020). This protein has a very high sequence similarity with the NMCP candidate identified in *P. margaritaceum,* however, both the N- and C-terminal domains are truncated relative to the bona fide candidate (pm001739g0030). Consequently, the shorter protein was discarded in our filtration protocol because it lacked significant intrinsically disordered regions at the N- and C-termini. Other top “hits” returned from the local BLAST searches had insignificant scores. Existing annotation and additional protein family analysis of these less significant “hits” revealed relationships to characterized proteins outside the nuclear lamina. These findings suggest that there is only one NMCP protein in *M. endlicherianum, Z. circumcarinatum, C. braunii,* and *K. nitens*, and possibly two in *P. margaritaceum*, but we found no evidence for NMCPs in *S. muscicola, C. atmosphyticus,* nor *M. viride*.

We next investigated whether these absences might result from genome misannotation. Using the 50-amino acid sequence of *P. patens,* we searched, using tBLASTn, both the transcriptomes and the genomes of three streptophytic algae taxa apparently lacking NMCPs. The transcriptome and genome scans of *C. atmosphyticus* and *M. viride* returned a few proteins in the search with marginally significant e-values; these proteins were the same as those returned in the BLAST search against the algal proteomes (see Supplementary Table 1). These results strengthen our initial conclusion that NMCP proteins are absent in *C. atmosphyticus,* and *M. viride*.

For *S. muscicola*, scans against the annotated transcriptome failed to return NMCP candidate proteins; however, tBLASTn searches against the genome identified three independent sequence scaffolds with putative exonic sequences capable of encoding a peptide with similarity to the conserved 50-amino acid region (see Supplementary Figure 1A). By analyzing RNA-seq reads aligned to genome scaffolds, we reconstructed the gene structure of three nearly complete NMCP genes in *S. muscicola* (see Supplementary Figure 1B and Supplementary File 1). The three deduced Spirogloea NMCP proteins share high identity (>80%) and contain a characteristic extended coiled-coil region and a C-terminal domain with a high intrinsic disorder score (Supplementary Figure 1C). Gaps in genome sequence obscure the true nature of the N-terminal domain, but each gene is expressed based on paralog-specific polymorphisms in the RNA-seq reads. A whole genome triplication was observed in *S. muscicola* (Cheng *et al*., 2019), consistent with our finding of three highly similar NMCP paralogs in this species.

We next examined the structural similarities among the four streptophytic algal NMCPs recovered by our filtration protocol. Although these proteins have a similar domain organization, they share only low amino acid sequence similarity, with the highest degree of sequence identity in the coiled-coil domain (21-29%) (Supplementary Figure 2). The four streptophytic algal proteins contain some of the amino acid motifs outside of the coiled-coil region that were originally identified by Ciska et al. (2019) as conserved among NMCP proteins in land plants and algae (Figure 5 and Supplementary Figure 3).

**Figure 5:**
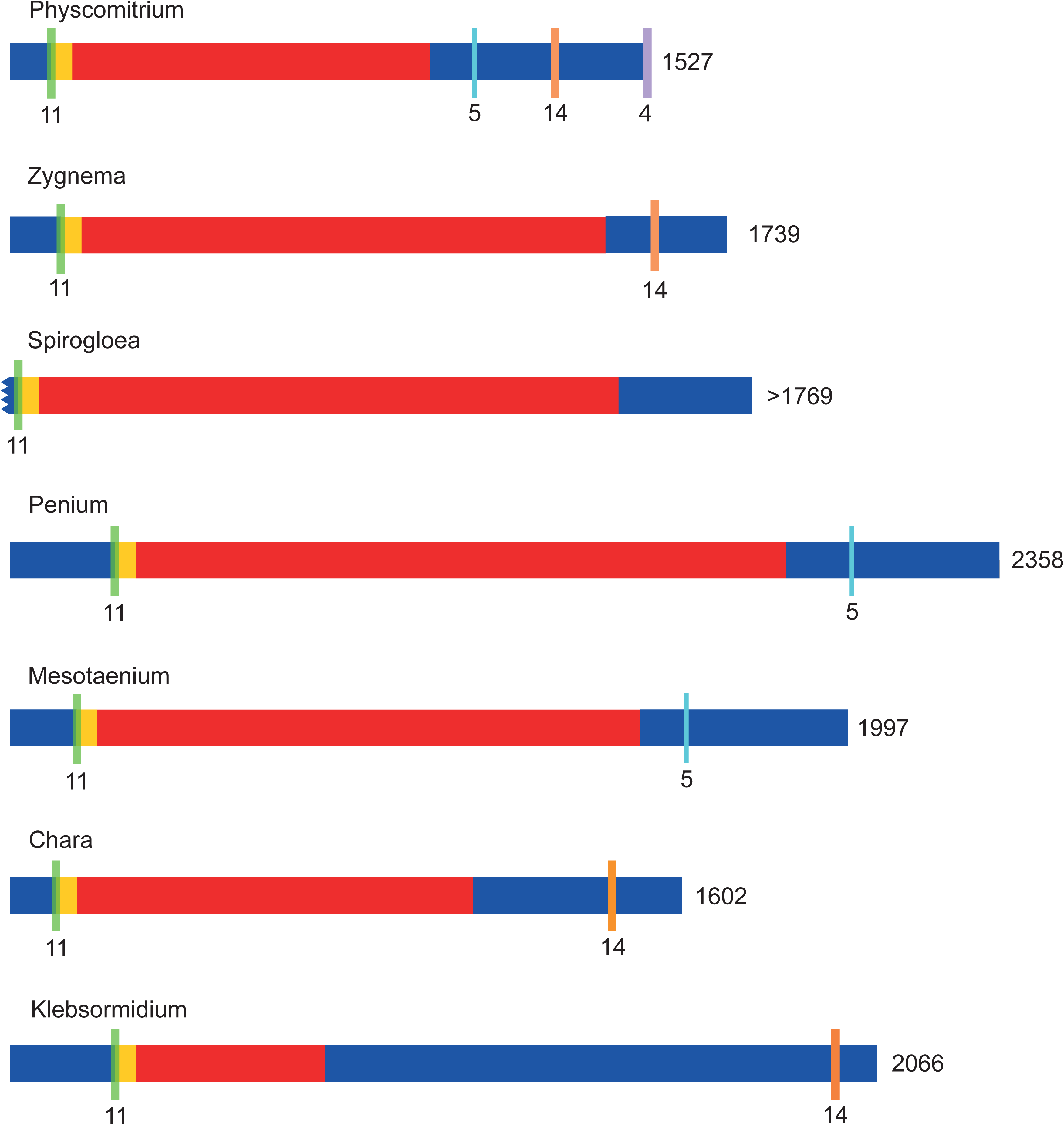
Conserved NMCP motifs across algal candidates. Algal candidate NMCPs and *P. patens* NMCP1 (A0A2K1L4A2_PHYPA) aligned showing the differences in domain size and presence of conserved regions identified in Ciska et al. (2019). Spirogloea is represented by NMCP A in this figure. Regions in blue represent intrinsic disordered domains, while regions in red represent the coiled-coil domain. The total length of each protein is indicated to the right, # of amino acids; domains are drawn to scale. The yellow box represents the 50-amino acid motif at the beginning of the coiled-coil domain described in this study (Figure 4). The head domain characterized as motif 11 in Ciska et al. (2019) is represented in green. The tail motif 5 is represented in aqua, motif 14 is represented in orange, and motif 4 is represented in purple (see Supplementary Figure 3).

The length of NMCP proteins and their constituent domains varies between angiosperms and streptophytic algae. Figure 5 illustrates a trend of increased size of NMCPs, particularly the intrinsically disordered region, moving from angiosperms to moss to streptophytic algae (Supplementary Figure 4). When the sizes of the three domains of the tripartite NMCPs were compared within and between land plants and streptophytic algae, we noted a larger variance among the algal proteins (see Figure 6). The size of the N-terminal domain averaged over 150 amino acids for the four algal proteins identified here, roughly twice the size of N-terminal domains of typical land plant NMCPs (Figures 5 and 6). A statistically significant difference was also seen in the length of the coiled-coil region, with the algal NMCPs having a longer central domain compared to land plant NMCPs. A similar trend was observed for the C-terminal region, although the differences in this case were not statistically significant (Figure 6). These comparisons demonstrate that considerable variation in NMCP domain size is tolerated in streptophytic algae while preserving the overall tripartite organization of the protein.

**Figure 6:**
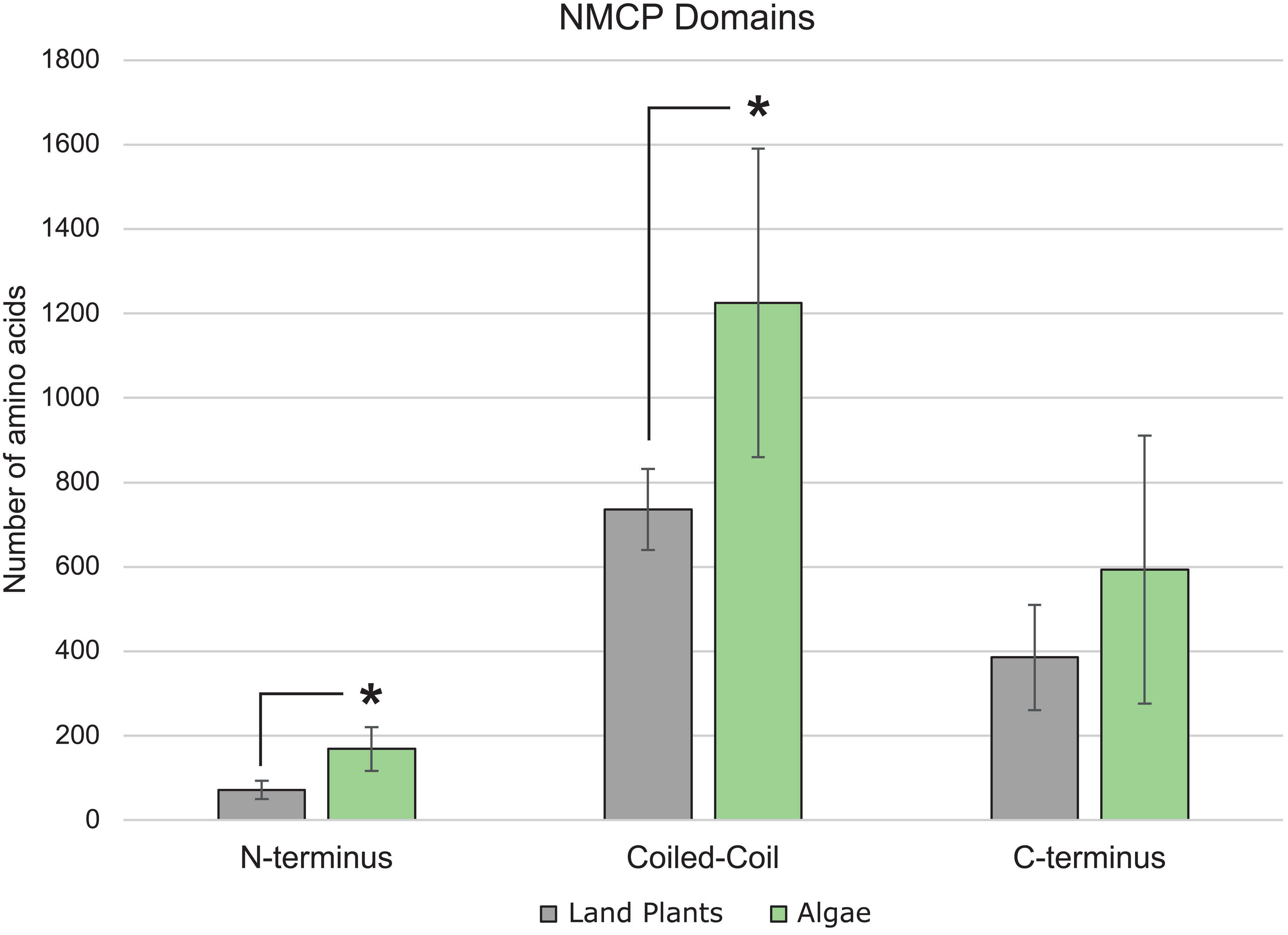
Comparison of the size of each tripartite domain in NMCPs between land plants and algae. Length of different domains (# of amino acids) of NMCP proteins from land plants compared to streptophytic algae. Representative proteins from land plants (n=8) are shown in gray, including *A. thaliana* CRWN1 and 4; *S. lycopersicum* NMCP1B and 2; *Salvinia cucullata* NMCP1 and 2; and both *P. patens* NMCPs. NMCP proteins from streptophytic algae (n=6) are shown in green and are composed of the identified NMCP candidates from the filtering protocol in *Z. circumcarinatum, S. muscicola*, *M. endlicherianum, C. braunii*, and *K. nitens*, and *P. margaritaceum*. Error bars indicate standard deviation. The asterisk over the algae N-terminus and coiled-coil domain indicates a significant difference in domain sizes assessed using a Tukey HSD test (p<0.01).

## DISCUSSION

NMCPs in land plants resemble lamins in structural organization, as illustrated in Figure 1. We reasoned that we might be able to identify transitional stages in the evolution of NMCPs by looking more closely for structural homologs in streptophytic algae in sister lineages of land plants. Would it be possible to start with validated NMCP-class proteins in land plants and bridge out through algae toward the Last Eukaryotic Common Ancestor to identify a connection to lamins? Or are NMCPs truly plant-specific (that is, limited to the ‘green lineage’)? If the latter, can we identify NMCPs in all lineages of photosynthetic organisms?

We developed a filtering protocol that considers the physical characteristics of proteins as criteria to search a proteome for potential homologs without dependence on sequence similarity. NMCP candidates could be identified in streptophytic algae through the analysis of four parameters: intrinsically disordered region content, peptide size, isoelectric point, and the presence of an extended coiled-coil domain. Candidate proteins were assessed by visualizing the organization of the tripartite structure with reference to known angiosperm NMCPs, such as *Arabidopsis* CRWN proteins. The filtration protocol (Figure 3) reduced the number of candidates to a small pool (Table 1), and we introduced a final step based on the presence of a weakly conserved region of 50 amino acids (Figure 4).

Through this process, we identified NMCP proteins in six streptophytic algal taxa: *Klebsormidium nitens*; *Chara braunii*; *Mesotaenium endlicherianum*; *Penium margaritaceum*; *Zygnema circumcarinatum*, and *Spirogloea muscicola* (Supplementary Table 2). The *Spirogleoa muscicola* genome, which underwent a recent triplication (Cheng *et al*., 2019), encodes three closely related NMCPs (Supplementary Figure 1 and Supplementary File 1), but the remaining four taxa contain only a single NMCP protein. Although we found a second NMCP paralog candidate in *Penium*, this protein contains only the coiled-coil region. It is possible that this second predicted *Penium* protein was derived from a full-length progenitor NMCP or that the unusual protein structure stems from misannotation of the gene.

Despite retention of a characteristic tripartite structure, the size of the domains varies considerably among the five algal NMCPs (Figure 5), and the overall size of the algal proteins tends to be larger than their counterparts in land plants (Figure 6). Further, the algal NMCPs share very limited amino acid similarity (Supplementary Figure 2), suggesting that primary sequence can diverge significantly if the proteins maintain the structural characteristics that define the NMCP family. Nonetheless, a few short sequence motifs shared among NMCP proteins can be identified among the algal proteins (see Figures 4 and 5) (Ciska *et al*., 2019). Absent, however, is a well-conserved C-terminal motif, which is present in many NMCP proteins in land plants and implicated in ABA regulation (Supplementary Figure 3) (Zhao *et al*., 2016). The function of the few short amino acid motifs shared by algal NMCPs is unknown; it is possible that these regions represent sites for interaction with core nucleoskeletal or nuclear membrane proteins.

A previous study by Koreny and Field (2016) uncovered NMCP candidates in streptophytic algae in the genera *Coleochaete, Klebsormidium, Nitella,* and possibly *Spirogyra*, and Ciska et al. (2019) added additional analysis. These NMCP proteins were identified using a similarity-based search from transcriptomic datasets, but the limitations of such an approach were acknowledged by Koreny and Field (2016), citing the extreme divergence of these proteins. In addition, similarity among even unrelated coiled-coil regions could lead to a high background of false positives (Koreny and Field, 2016). We examined the three novel algal NMCP candidates identified by Koreny and Field (2016) using the structural characteristics described in this study (note that the previous study also found the protein from *Klebsormidium* uncovered in our filtration protocol). Two of the three NMCP candidates conform to our structural criteria (JV767595 from *Nitella mirabilis*; and GB-SL01031242 from *Coleochaete orbicularis*); however, the *Spirogyra* protein (GB-SM01021289 from *Spirogyra pratensis*) is a relatively short protein (634 amino acids) that lacks significant intrinsically disordered regions.

While NMCPs are clearly present in different streptophytic algae, these nuclear lamina proteins are not found in all streptophytes. We conclude that NMCPs are absent from the proteomes of *Chlorokybus melkonianii* and *Mesostigma viride* based on the following results. First, we analyzed the proteins in our finalist candidate list shown in Table 1 for the possibility that they might represent previously unrecognized NMCP variants. However, these candidates were related to proteins with roles outside the nuclear lamina and therefore are unlikely to be highly diverged NMCP homologs. Second, to identify homologs that might not have the diagnostic tripartite domain organization, we searched the seven complete proteomes using a query corresponding to a 50-amino acid block in the coiled-coil region from a *Physcomitrium patens* NMCP (Figure 4). No new NMCP candidates were recovered by this approach from proteomes of *Chlorokybus melkonianii* or *Mesostimga viride*. We did, however, identify an additional NMCP-like protein from *Penium margaritaceum* that eluded the filtering protocol because it lacked disordered domains as noted above. Third, to account for NMCP proteins potentially overlooked due to errors in genome annotation, we performed tBLASTn searches with the same query against algal transcriptomes, as well as genome sequences, and found no new candidates. These findings argue that *Chlorokybus melkonianii* and *Mesostimga viride* lack NMCP proteins.

Mapping the presence and absence of NMCP proteins relative to the phylogeny of streptophytic algae demonstrates that NMCPs do not appear at the boundary between green algae (Chlorophyta) and streptophytic algae. The streptophytes most closely related to green algae, *Mesostigma viride* and *Chlorokybus melkonianii,* do not have a recognizable NMCP. Moving toward lineages more closely related to land plants, NMCPs first appear in Klebsormidophyceae, represented by *Klebsormidium nitens*. Taking a further step toward land plants, NMCPs are present in the Charophyceae. Within the Zygnematophyceae, we found NMCPs in all three different taxa examined. These data indicate that NMCP proteins were first acquired in the green lineage at the branch point distinguishing the extant Classes Mesostigmatophyceae and Klebsormidophyceae (Figure 7). Our results, which demonstrate that NMCP-class nuclear lamina proteins are present in some but not all taxa in the green lineage, are consistent with the convergent evolution model for the nuclear lamina in streptophytes. We found no evidence for intermediate NMCP variants, resembling ancestral lamins, corresponding to more basal nodes of the green lineage. Thus, we conclude that NMCPs are plant-specific proteins with an independent origin and are not highly diverged lamins.

**Figure 7:**
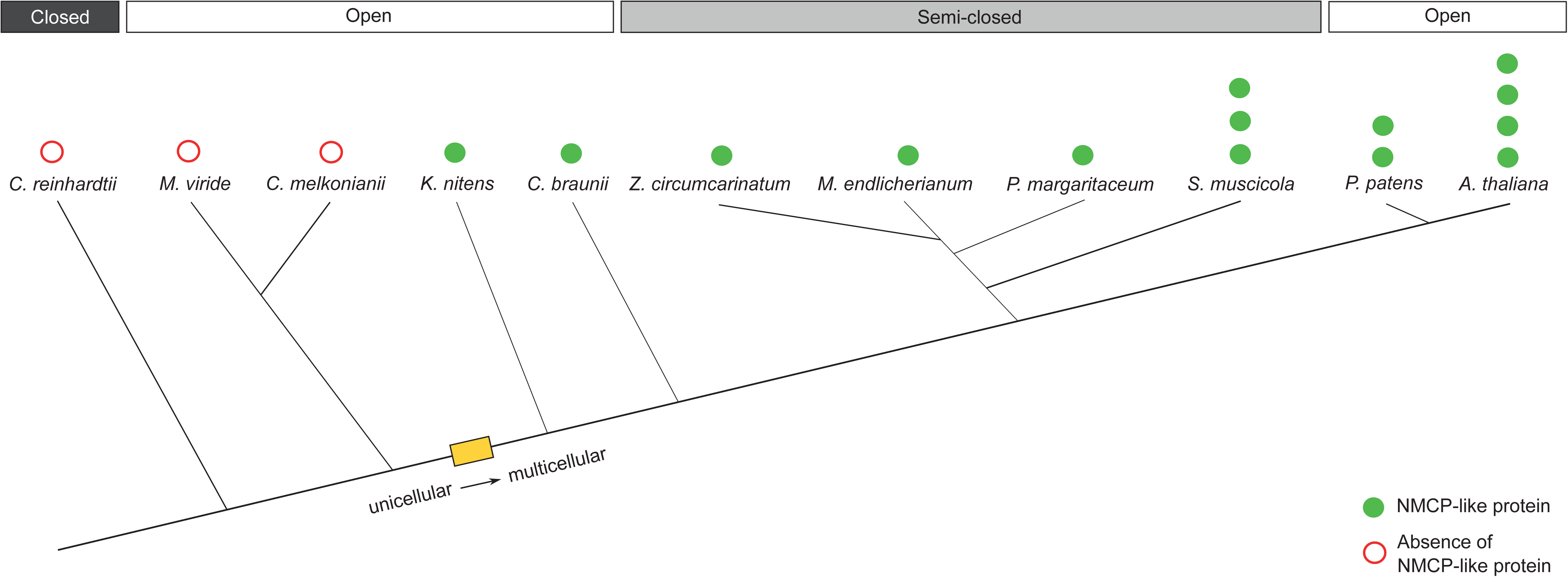
Distribution of NMCP proteins in the green lineage. Presence or absence of NMCP or NMCP-like protein by species and number of proteins identified. Branch lengths are proportional to the tree shown in Figure 2. The distribution of mitotic types is shown above the tree. The position of the transition between unicellularity and multicellularity is indicated by the yellow rectangle (Umen, 2014; but see Bierenbroodspot *et al*., 2024).

The distribution of NMCP proteins in streptophytic algae provides the opportunity to test the hypothesis that NMCPs play a role in the control of nuclear dynamics disassembly and reassembly during mitosis, analogous to lamins (Goldman *et al*., 2002). According to this hypothesis, NMCPs would be required to carry out an open mitosis, such as that which occurs in land plants, where the nuclear envelope breaks down before chromosomal segregation and reassembles after division (Rose, 2007). In closed mitosis, the nuclear envelope remains intact throughout division, and the nucleus partitions into two daughter nuclei during cytokinesis (Heath, 1986). Consistent with the hypothesis, green algae (Chlorophyta), which undergo closed mitosis, lack NMCPs (Heath, 1986; Koreny and Field, 2016). Bracketed by green algae and land plants, the streptophytic algae display a range of different mitotic types, including open, semi-closed, and closed. In some cases, streptophytes that contain NMCPs, including in the genus *Klebsormidium, Nitella*, and *Chara*, exhibit an open mitosis (Heath, 1986). The Zygnematophyceae have a semi-closed mitosis, characterized by partial disassembly of the nuclear envelope (Van Den Hoek *et al*., 1995). This consideration suggests that all taxa in Zygnematophyceae should contain an NMCP to mediate a partial nuclear breakdown, and our results correspond to this prediction (Figure 7). The state of the nucleus during mitotic division, however, is not necessarily predicted by presence or absence of an NMCP homolog. In certain streptophytes more distantly related to land plants, open mitosis occurs without NMCPs. *Mesostigma* is believed to be the final genus included in the Streptophyta at the boundary between the Chlorophyta (Rodríguez-Ezpeleta *et al*., 2007). Its sister group, represented in our study by *Chlorokybus melkonianii*, has an open mitosis but does not have an NMCP (Lokhorst *et al.,* 1988). Therefore, no simple correlation exists between mitotic type and the presence/absence of NMCPs in streptophytic algae, arguing against the hypothesis that acquisition of NMCPs enabled open mitosis in the green lineage. The specific roles played by NMCPs in nuclear dynamics during mitosis remain to be defined, but we anticipate that investigations exploiting the diversity of nuclear types and nuclear lamina components in streptophytic algae will help elucidate these roles.

It is notable that the position on the phylogenetic tree corresponding to the acquisition of NMCPs coincides with the transition from unicellularity to multicellularity in the streptophytic algae (see Figure 7). This finding suggests that the nuclear lamina might be required to complete the mechanical integration among nuclei of different cells in a single organism (Maurer and Lammerding, 2019), mediated via the cytoskeleton and LINC (LINKER OF NUCLEOSKELETON AND CYTOSKELETON) complexes (Chang *et al*., 2015; Meier *et al*., 2017; Wong *et al*., 2021)., Alternatively, NMCP proteins and a nuclear lamina might be needed to buffer more substantial mechanical stresses exerted on the nucleus through the cytoskeleton due to external forces acting on the surface of a larger organism or originating from physical strain between attached cells.

## MATERIALS AND METHODS

Each proteomic filtration consisted of four general steps, compounding the previous results before advancing. Four physical characteristics were assessed for filtration: Amino acid chain length, intrinsically disordered regions, isoelectric point, and coiled-coil patterns. In Arabidopsis, CRWN proteins are well studied and provide a convenient positive control and a point of reference for the filtration methods. For example, the CRWN1 protein is 1132 amino acids long, and a threshold of 800 was chosen to account for potential candidates with fewer amino acids than CRWN1. The coiled-coil region of CRWN1 is about 700 amino acids long and has a short N-terminus of about 50 amino acids.

Metapredict, a machine learning-based intrinsic disorder prediction software, generates IDR predictions for each protein from the amino acid sequence (Meier *et al*., 2017). For all proteomic datasets, IDRs were predicted for each individual protein, and values for angiosperms and gymnosperms were compared to the existing literature to confirm their accuracy. Metapredict returned disorder probability scores describing the confidence of a disordered region on a per residue basis across each protein for a complete proteome. These scores were consolidated to a per protein basis by converting per residue scores to a binary system. The developers of Metapredict recommend that a score of 0.3 constitutes significant disorder; subsequently, the threshold of the binary systems was set at 0.3. Taking the binary results per residue, each protein was assigned a percentage of residues satisfying the threshold. The output reported the number of total residues, as well as the number of disordered amino acids and total number of amino acids, allowing for amino acid chain length to be filtered simultaneously with disorder.

The proteins’ isoelectric points were calculated in parallel with filtration for overall size and disorder content. The isoelectric point (IEP) of a protein, or pKi, describes the average pKa of each amino acid residue over an entire protein. The IEP allows for the visualization of a protein’s optimal conditions to function. Lamin isoelectric points are typically acidic, ranging from about 4-6 pKi (Krohne and Benavente, 1986). The bioinformatics package EMBOSS was used for local isoelectric point prediction in UNIX (Rice *et al*., 2000). Each protein returned a single score which was then filtered to satisfy the acidic range of lamin pKi.

Filtered lists of proteins for size, IDR, and IEP were compiled and consolidated using Venny’s biological Venn diagram tool to select for proteins that satisfied all three criteria (https://bioinfogp.cnb.csic.es/tools/venny/). A FASTA file of only the selected proteins was then generated and filtered for the coiled-coil domain.

The presence of a long coiled-coil domain is strongly characteristic of lamin proteins. To model these regions computationally, the program DeepCoil, a coiled-coil domain prediction program, was used to assign each residue a probability of being part of a coiled-coil in the tertiary structure (Ludwiczak *et al*., 2019). The scores of each protein were visualized graphically and analyzed for similar patterns to the coiled-coil domain of CRWN1 in *A. thaliana.* A long central coiled-coil, with a disordered N-terminus of about 100 amino acids and a disordered C-terminus of varying length is indicative of an NMCP-like protein. Each coiled-coil output was assessed visually for these characteristics and was assigned a similarity grouping from 1 (no coiled-coil) to 5 (very similar coiled-coil pattern). Typically, only proteins with a score of 5 were advanced to the final step of the analysis in which the coiled-coil and disorder plots were overlayed.

## ACKNOWLEDGEMENTS

We are grateful for the work of Rachel Christopherson, who worked on an early version of this project. The authors thank the anonymous reviewer of the manuscript for their extremely helpful suggestions and insights. We also thank Adrian Powell, Ryan Preble, and other members of the Boyce Thompson Institute Computational Biology Center for their technical support and advice. This work was supported by funding from the National Science Foundation [grant URoL-2022048 to EJR].

## Data Availability

Genome and Proteome Sources

The *Arabidopsis thaliana* proteome was obtained via TAIR (www.arabidopsis.org). The proteome for *Penium margariticeum* was obtained from the Zhangjun Fei lab (Jiao *et al*., 2020). The proteome for *Klebsormidium nitens* (Hori et al 2014), *Chlamydomonas reinhardtii* (Merchant *et al*., 2007), and *Physcomitrium patens* (Rensing *et al*., 2008) were obtained via the NCBI’s GenBank genome portal. The proteomic information for the remaining algae: *Zygnema circumcarinatum* (Feng *et al*., 2024), *Mesotaenium endlicherianum* (Dadras *et al*., 2023)*, Spirogloea musicola* (Cheng *et al*., 2019)*, Mesostigma viride* (Wang *et al*., 2020)*, Chlorokybus melkonianii* (Wang *et al*., 2020), and *Chara braunii* (Nishiyama *et al*., 2018) were downloaded from the JGI genome portal (https://genome.jgi.doe.gov/portal/). Links to each dataset are provided in Supplementary Table 3.

## Code Availability

The code used for the filtration pipeline in this manuscript can be accessed at https://github.com/brendankz/Proteome-filtration.

## Notes

### Competing Interest Statement

The authors have declared no competing interest.

### Summary of Updates

The manuscript and the supplemental files have been updated and include additional results and analysis.

