## Supplemental Information for "Structural diversity and distribution of NMCP-class nuclear lamina proteins in streptophytic algae"

### SUPPLEMENTARY INFORMATION

**Supplementary Table 1.** BLAST results using the *P. patens* NMCP1 coiled-coil 50-amino acid region as a query

| Species | E-Value | Accession/Locus | Annotation |
|---|---|---|---|
| *Mesotaenium endlicherianum* | 9.00E-06 | ME000069S09116 | NMCP Candidate |
| | 1.1 | ME000031S05348 | RNA binding, helicase-like |
| | 1.9 | ME000288S04886 | NET actin-binding domain protein |
| | 1.9 | ME000288S04885 | NET actin-binding domain protein |
| | 2.3 | ME000276S04628 | Coiled-coil domain protein |
| *Penium margaritaceum* | 5.00E-05 | pm001739g0030 | NMCP Candidate |
| | 0.001 | pm015243g0010 | Putative nuclear matrix constituent protein 1-like |
| | 1.3 | pm002294g0040 | Pre-mRNA-splicing factor 38B |
| | 3.4 | pm027102g0020 | Pre-mRNA-splicing factor 38B |
| | 5.3 | pm056296g0010 | Pre-mRNA-splicing factor 38B |
| *Spirogloea muscicola* | 0.2 | SM000034S12656 | Myosin heavy chain-like protein |
| | 0.23 | SM000112S23966 | Myosin heavy chain-like protein |
| | 1.2 | SM000205S06221 | Myosin heavy chain-like protein |
| | 2.1 | SM000082S22847 | Aminotransferase-like |
| | 2.4 | SM000057S18360 | Aminotransferase-like |
| *Chara braunii* | 2.00E-04 | Chabra1336100rna-CBR_g27897 | NMCP Candidate |
| | 0.52 | Chabra1347117rna-CBR_g41659 | HSP70/HSP90 binding |
| | 0.72 | Chabra1342418rna-CBR_g37091 | Ubiquitin ligase |
| | 3.4 | Chabra1324201rna-CBR_g58401 | WD40/YVTN repeat domain superfamily |
| | 3.7 | Chabra1337629rna-CBR_g30118 | STE20-related kinase adapter protein alpha/beta-like |
| *Klebsormidium nitens* | 2.00E-05 | GAQ84530.1 | NMCP Candidate |
| | 0.67 | GAQ90068.1 | Coiled-coil domain protein |
| | 0.71 | GAQ78086.1 | Calponin-homology domain |
| | 1.7 | GAQ91089.1 | Coiled-coil domain protein |
| | 2 | GAQ86916.1 | Beta-tubulin folding cofactor D |
| *Mesostigma viride* | 0.12 | Mv13079-RA.1 | WD40 repeat domain superfamily |
| | 0.2 | Mv11449-RA.3 | Centrosomal Protein 2 |
| | 1.2 | Mv10613-RA.1 | Zinc finger, RING type |
| | 1.6 | Mv02123-RA.2 | Centrosomal Protein 2 |
| | 1.6 | Mv14161-RA.1 | Kelch propeller motif |
| *Chlorokybus melkonianii* | 1.6 | Chrsp134S02085 | Kinesin-like protein |
| | 2.2 | Chrsp134S02085 | Kinesin-like protein |
| | 2.4 | Chrsp26S00300 | DUF5695 |
| | 3.3 | Chrsp57S06825 | NADH-dependent oxidoreductase modular protein |
| | 3.4 | Chrsp17S02814 | Malonyl-CoA decarboxylase |
| *Zygnema circumcarinatum* | 4.90E-04 | UTEX1559_01932.1.v1 UTEX1559_10991.1 | NMCP Candidate |
| | 0.59 | UTEX1559_10850.1 | Myosin-like |
| | 0.59 | UTEX1559_10503.1.v1 | Mysosin-like |
| | 3.2 | UTEX1559_05347.1 | Beta-arabinofuranosyltransferase RAY1-like |

The 50-amino acid coiled-coil motif from *P. patens* NMCP1 was used as a BLAST query against the proteome (BLASTp) and transcriptome (tBLASTn) of the eight algal species considered in this study. The results from each search and their significance are shown in the table above. Five of the six NMCPs (highlighted in blue) were recovered with the *P. patens* query. The *Spirogloea* NMCPs were not recovered as these proteins and genes were not included in the proteome nor the annotated transcriptome dataset. Other proteins returned were not statistically significant matches, and most mapped to proteins with predicted roles outside the nucleus. The two NMCP candidates recovered from *Z. circumcarinatum* are encoded from the same location in the genome, and represent minor annotation variants of the same protein, despite the different locus designations.

**Supplementary Table 2.** Algal NMCP candidates and reference land plant NMCPs

| Protein ID | Species | Disorder | Size | Iso-electric Point |
|---|---|---|---|---|
| CRWN1 | *Arabidopsis thaliana* | 0.4417 | 1132 | 4.9613 |
| CRWN4 | *Arabidopsis thaliana* | 0.3218 | 1010 | 4.8438 |
| NMCP1, A0A2K1L4A2_PHYPA | *Physcomitrium patens* | 0.4768 | 1527 | 4.5084 |
| NMCP2, A0A2K1L6J7_PHYPA | *Physcomitrium patens* | 0.501 | 1549 | 4.3949 |
| NMCP A Contig s_61 | *Spirogloea muscicola* | 0.444 | 1769 | 4.9801 |
| NMCP B Contig s_257 | *Spirogloea muscicola* | 0.437 | 1768 | 5.0449 |
| NMCP C Contig s_265 | *Spirogloea muscicola* | 0.452 | 1765 | 4.9722 |
| UTEX1559_01932.1.v1 | *Zygnema circumcarinatum* | 0.394 | 1739 | 5.8043 |
| ME000069S09116 | *Mesotaenium endlicherianum* | 0.677 | 1352 | 4.6971 |
| pm001739g0030 | *Penium margaritaceum* | 0.5704 | 1345 | 4.8151 |
| Chabra1336100rna-CBR_g27897 | *Chara braunii* | 0.52 | 1602 | 5.0675 |
| A0A1Y1I0R3_KLENI | *Klebsormidium nitens* | 0.8432 | 2066 | 4.8661 |

Values for land plant and algal NMCP's predicted intrinsic disorder, isoelectric point, and number of amino acids. Disorder scores are probability values on a scale of 0 to 1, size is represented by the number of amino acids, and iso-electric point represents pKi.

**Supplementary Table 3.** Proteome file sources

| **Species** | **Proteome Source** |
|---|---|
| *Arabidopsis thaliana* | https://www.arabidopsis.org/download/index-auto.jsp?dir=/download_files/Proteins |
| *Physcomitrium patens* | https://www.ncbi.nlm.nih.gov/datasets/taxonomy/3218 |
| *Penium margaritaceum* | http://bioinfo.bti.cornell.edu/Penium/ |
| *Mesotaenium endlicherianum* | https://phycocosm.jgi.doe.gov/Mesen1_1/Mesen1_1.home.html |
| *Zygnema circumcarinatum* | https://genome.jgi.doe.gov/portal/pages/dynamicOrganismDownload.jsf?organism=Zygcir1559_1 |
| *Spirogloea muscicola* | https://phycocosm.jgi.doe.gov/Spimu1_1/Spimu1_1.home.html |
| *Chara braunii* | https://phycocosm.jgi.doe.gov/Chabra1/Chabra1.home.html |
| *Klebsormidium nitens* | https://www.ncbi.nlm.nih.gov/datasets/genome/?taxon=105231 |
| *Mesostigma viride* | https://phycocosm.jgi.doe.gov/Mesovir1_1/Mesovir1_1.home.html |
| *Chlorokybus melkonianii* | https://phycocosm.jgi.doe.gov/Chlat1_1/Chlat1_1.home.html |
| *Chlamydomonas reinhardtii* | https://www.ncbi.nlm.nih.gov/datasets/genome/GCF_000002595.2/ |

Link to proteome source by species.

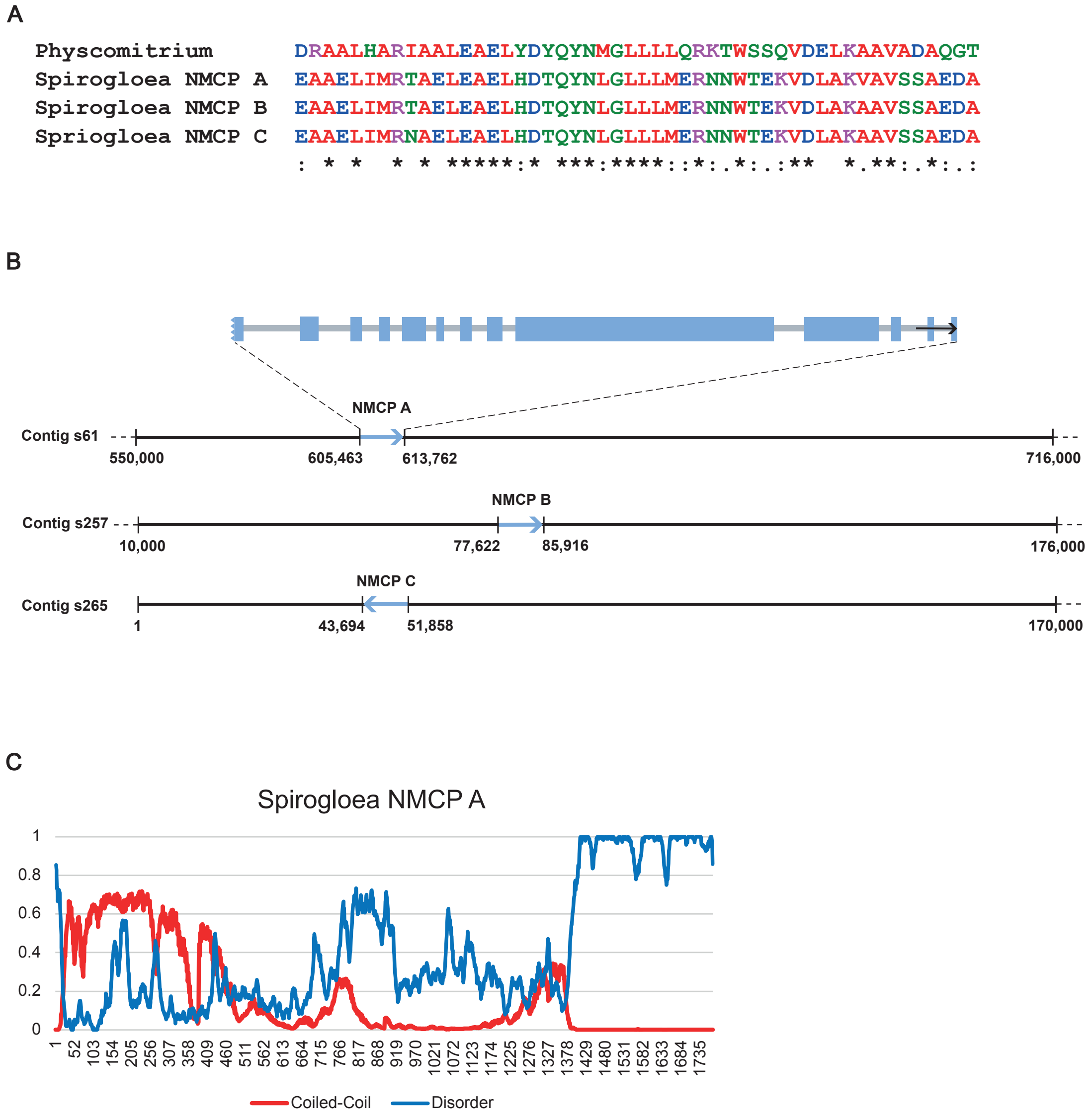


**Supplementary Figure 1.** The *Spirogloea muscicola* genome encodes three NMCP proteins

(a) Multiple sequence alignment between *Physcomitrium patens* NMCP1 (A0A2K1L4A2) and the three *Spirogloea* NMCP, designated A, B, and C, in the 50-amino acid region of the coiled-coil domain identified in Figure 4.

(b) Intron/exon structure for *Spirogloea* NMCP A. Exons are represented as light blue rectangles; the potentially truncated N-terminus is denoted by a jagged edge on exon 1. The locus of each NMCP gene is shown below, identifying the sequence in the corresponding scaffolds provided by the available sequencing data for *Spirogloea muscicola* (Cheng *et al.*, 2019).

(c) Coiled-coil and intrinsically disordered domain scores for *Spirogloea* NMCP A. The red line shows the probability of each amino acid being part of the coiled-coil domain, while the blue line represented the probability that an amino acid is intrinsically disordered.

(A) N-terminal domain

| | *Physcomitrium* | *Spirogloea* | *Penium* | *Mesotaenium* | *Zygnema* | *Chara* | *Klebsormidium* |
|---|---|---|---|---|---|---|---|
| *Klebsormidium* | 20 | | 17 | 23 | 19 | 16 | 100 |
| *Chara* | 10 | | 6 | 15 | 18 | 100 | |
| *Zygnema* | 21 | | 15 | 22 | 100 | | |
| *Mesotaenium* | 19 | | 17 | 100 | | | |
| *Penium* | 16 | | 100 | | | | |
| *Spirogloea* | | 100 | | | | | |
| *Physcomitrium* | 100 | | | | | | |

(B) Coiled-coil domain

| | *Physcomitrium* | *Spirogloea* | *Penium* | *Mesotaenium* | *Zygnema* | *Chara* | *Klebsormidium* |
|---|---|---|---|---|---|---|---|
| *Klebsormidium* | 21 | 19 | 25 | 24 | 20 | 22 | 100 |
| *Chara* | 25 | 23 | 24 | 26 | 23 | 100 | |
| *Zygnema* | 23 | 19 | 25 | 24 | 100 | | |
| *Mesotaenium* | 28 | 23 | 29 | 100 | | | |
| *Penium* | 25 | 25 | 100 | | | | |
| *Spirogloea* | 26 | 100 | | | | | |
| *Physcomitrium* | 100 | | | | | | |

(C) C-terminal domain

| | *Physcomitrium* | *Spirogloea* | *Penium* | *Mesotaenium* | *Zygnema* | *Chara* | *Klebsormidium* |
|---|---|---|---|---|---|---|---|
| *Klebsormidium* | 14 | 16 | 15 | 20 | 18 | 17 | 100 |
| *Chara* | 15 | 16 | 18 | 14 | 16 | 100 | |
| *Zygnema* | 14 | 18 | 18 | 16 | 100 | | |
| *Mesotaenium* | 14 | 15 | 18 | 100 | | | |
| *Penium* | 15 | 15 | 100 | | | | |
| *Spirogloea* | 15 | 100 | | | | | |
| *Physcomitrium* | 100 | | | | | | |

| | | | | | |
|---|---|---|---|---|---|
| 0-4% | 5-9% | 10-14% | 15-19% | 20-24% | 25-29% |

**Supplementary Figure 2.** Similarity among algal NMCP proteins. Pairwise analysis of amino acid identity, broken down by domain (N-terminal, coiled-coil, and C-terminal) among streptophytic NMCPs and Physcomitrium's NMCP1 (A0A2K1L4A2). Protein domains were defined by the predicted presence of a coiled-coil domain and corresponded to the domains represented in Figure 6. Due to the uncertainty and likely incomplete nature of the Spriogloea NMCP N-terminal regions, a *Spirogloea* NMCP representative was omitted from the N-terminal identity comparisons. The identity values among protein domains are represented by a heat map starting from light yellow, indicating low identity, ranging to dark red, indicating higher relative identity. The site CLUSTAL W tool on the Genome.jp site (https://www.genome.jp/tools-bin/clustalw) was used to generate identity scores.

```
Conserved Domain 11 GGLDEESLERKDRAALJAYI

Physcomitrella      GALDISSLERKDRAALHARI
Spirogloea          GALDEAGAARAEAAELIMRT
Penium              NWLDEVTMLRKERDELMLKV
Mesotaenium         GTLDEDSVVRKERGALQARV
Chara               GALSEVGLEKKERNFMVDHI
Klebsormidium       GALDEEVLVRREKEELQRQL
Zygnema             GSFDENTILRDEIDALNARV
                    . *.     : :  :  :
```

```
Conserved Domain 5 SWL-RKCASKIF

Physcomitrella     AWI-QRCASRAS
Penium             GWFMERCRRLL-
Mesotaenium        RWVLDRCSQLF-
                    *. :**
```

```
Conserved Domain 14 QTPGEKRYNLRRSTIVNTVA

Physcomitirum       GTPATKRYNFRPTTIVNMMG
Chara               KRERTVKYNLRRTTVYHQAK
Klebsormidium       MSAGLRRYNLRKSTLVKMHI
Zygnema             RPGGIQTQTIREVEVVNEVR
                            .:*   : :
```

```
Conserved Domain 4 EDEEPGEASIGKKLWNFLTT

Physcomitrella     PDDEGPTPTIREKIWDFLTT
```

**Supplementary Figure 3.** Conserved NMCP motifs. Multiple sequence alignments of algal NMCP proteins and *P. patens* NMCP1 of four conserved domains, excluding coiled-coil regions, identified in Ciska et. al (2019) and shown in Figure 5. Domain 11 corresponds to a conserved region in the head domain, domains 4, 5, and 14, are conserved sequences located in the C-terminus of NMCPs. Amino acid colors scheme is described in the legend in Figure 4. The symbols under the alignments reflect the similarites (. < :) or identities (*) among the sequences shown in colored text.

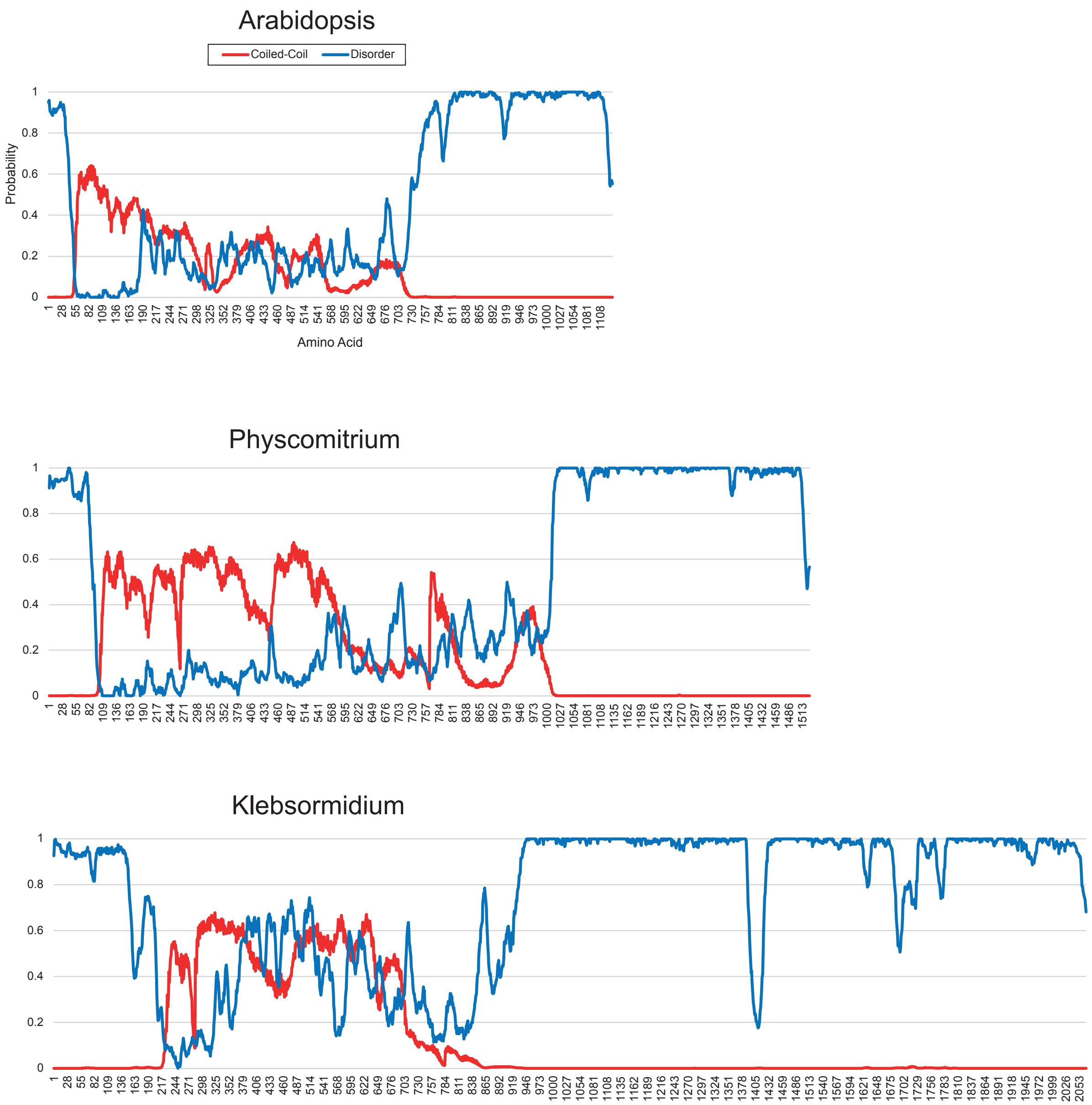


**Supplementary Figure 4.** Structural comparison of select NMCP proteins. Scores for intrinsically disordered (blue) and coiled-coil (red) regions were mapped per residue for NMCP proteins from *A. thaliana* (CRWN1), *P. patens* NMCP1 (A0A2K1L4A2_PHYPA), and *K. nitens* aligned by protein size. The x-axis shows the amino acid residue and the y-axis shows the probability of that residue being a part of a coiled-coil or disordered domain.

**Supplementary File 1.** Sequences of deduced *Spirogloea* NMCP proteins in fasta format
